## Supporting Information for "Integrating Self-Attention Transformer with Triplet Neural Networks for Protein Gene Ontology Prediction"

**Supplementary Materials**

**Table of content**

**Supporting Figures**

- Figure S1. The mean AUROC values of GO terms versus twelve GO prediction methods in four ranges. **(a)** range 5-10. **(b)** range 10-30. **(c)** range 30-50. **(d)** range >50.
- Figure S2. The median AUROC values of GO terms versus twelve GO prediction methods in four ranges. **(a)** range 5-10. **(b)** range 10-30. **(c)** range 30-50. **(d)** range >50.
- Figure S3. The distributions of AUROC values for GO terms in two ranges versus twelve GO prediction methods, where the median line in the box is the median AUROC value. **(a)** range 30-50. **(b)** range >50.
- Figure S4. The AUPR values of ten GO prediction methods under the sequence identity cut-off $t_{1}=30\%$ for three GO aspects on four individual species in CAFA3 test dataset. **(a)** Human **(b)** Arabidopsis **(c)** Fission Yeast **(d)** Mouse.
- Figure S5. The architectures of different models in ablation study.

**Supporting Tables**

- Table S1. The prediction performance with including root GO terms for ATGO and TALE on all 1068 test proteins. *p*-values in parenthesis are calculated between ATGO and TALE by one-tailed Student’s t-test. Bold fonts highlight the best performer in each category.
- Table S2. The summary of the proposed ATGO/ATGO+ and other ten competing GO prediction methods on a subset of 562 test proteins which have available templates or interaction partners in all of SAGP, PPIGP, FunFams, and DIAMONDScore. *p*-values in parenthesis are calculated between ATGO and other single-based methods and between ATGO+ and other composite methods by one-tailed Student’s t-test. Bold fonts highlight the best performer in each category.
- Table S3. The performance of ten GO prediction methods under the cut-off $t_{1}=30\%$ on 1177 no-knowledge (NK) and 2151 limited-knowledge (LK) CAFA3 proteins. Bold fonts highlight the best performer in each category.
- Table S4. The numbers of proteins for 20 species in CAFA3 test dataset.
- Table S5. The prediction performance of five GO prediction methods under the cut-off $t_{1}=100\%$ on CAFA3 test proteins. Bold fonts highlight the best performer in each category.
- Table S6. The incorrectly predicted GO terms for twelve methods on three proteins in BP aspect.
- Table S7. The numbers of proteins and GO terms in benchmark dataset.
- Table S8. The values of $margin$, $c_{f}$, $\alpha$, and $K$ for three GO aspects.

**Supporting Texts**

- Text S1. Sequence alignment-based GO prediction.
- Text S2. Protein-protein interaction-based GO prediction.
- Text S3. Naïve-based GO prediction.
- Text S4. The mathematics formulas for ESM-1b transformer.
- Text S5. The functional similarity between two proteins.

**Supporting Figures**

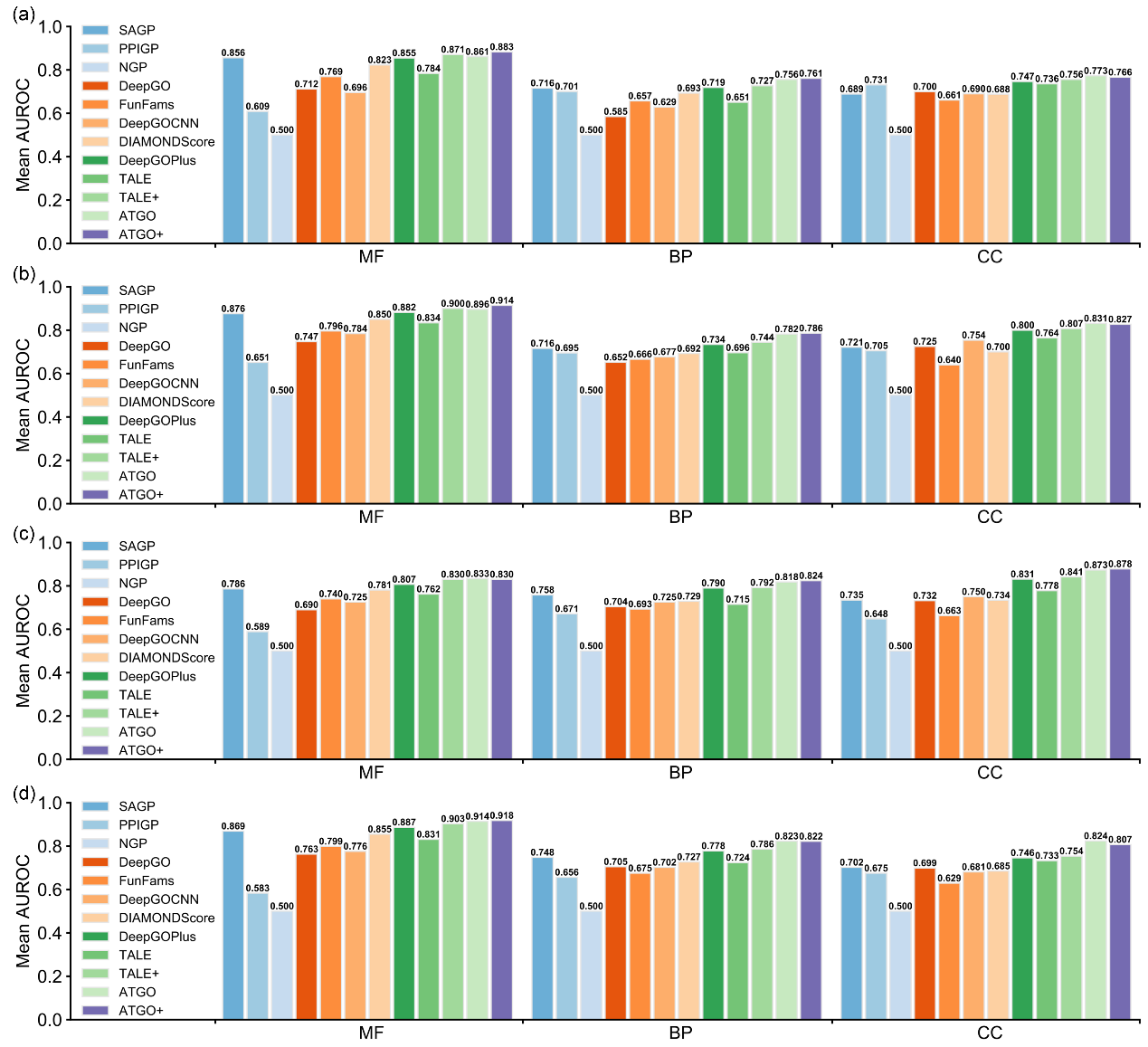

**Fig S1.** The mean AUROC values of GO terms versus twelve GO prediction methods in four ranges. **(a)** range 5-10. **(b)** range 10-30. **(c)** range 30-50. **(d)** range >50.

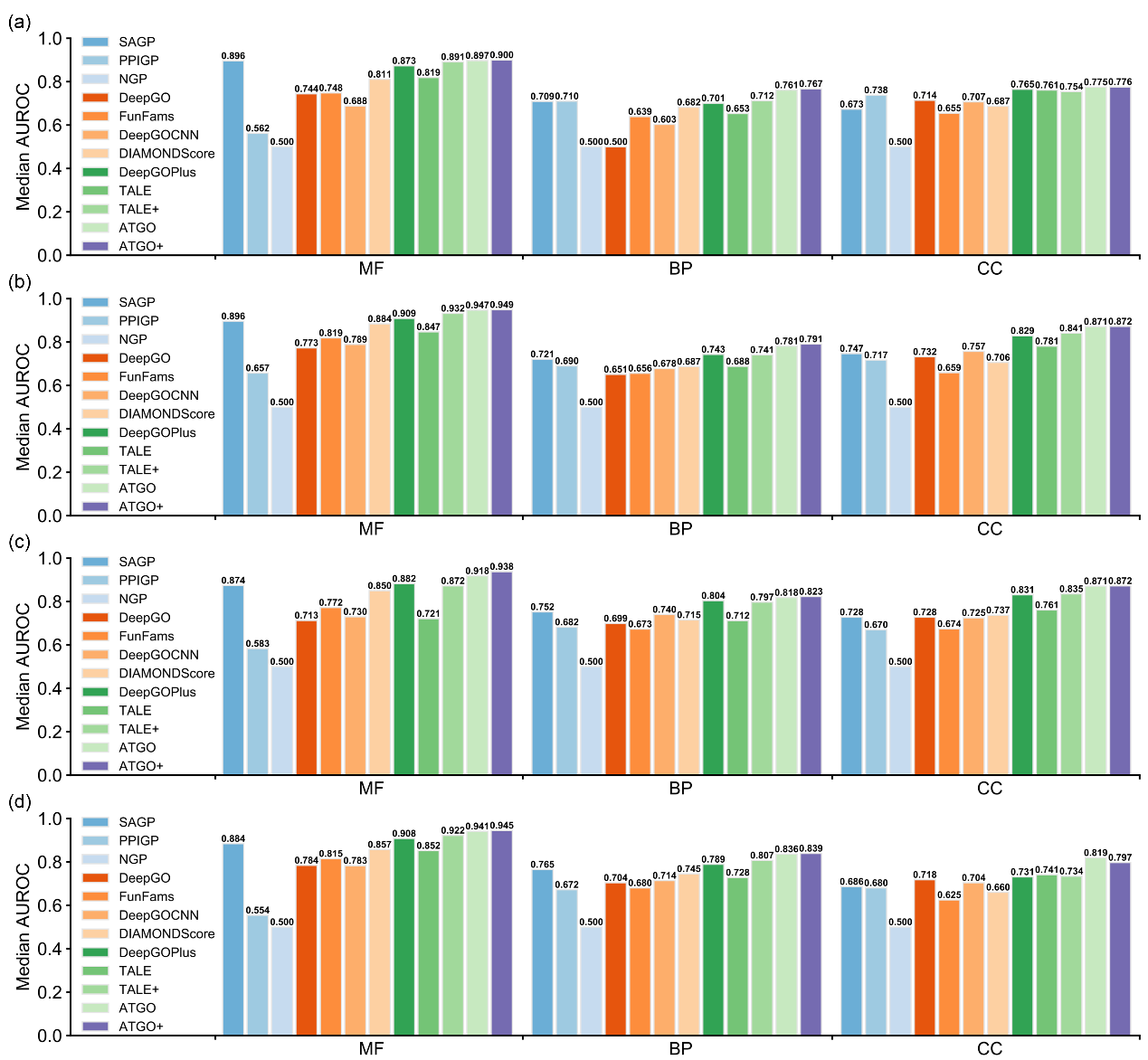

**Fig S2.** The median AUROC values of GO terms versus twelve GO prediction methods in four ranges. **(a)** range 5-10. **(b)** range 10-30. **(c)** range 30-50. **(d)** range >50.

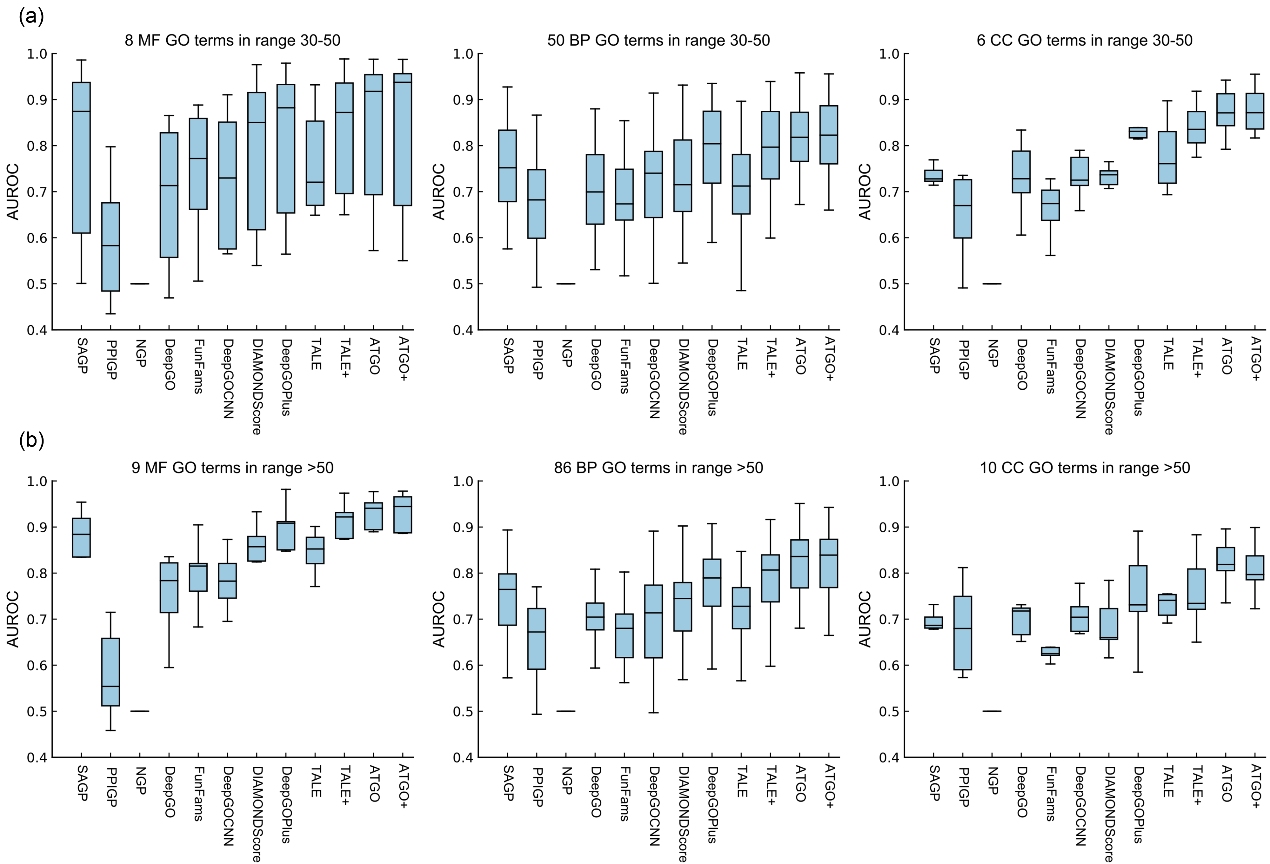

**Fig S3.** The distributions of AUROC values for GO terms in two ranges versus twelve GO prediction methods, where the median line in the box is the median AUROC value. **(a)** range 30-50. **(b)** range >50.

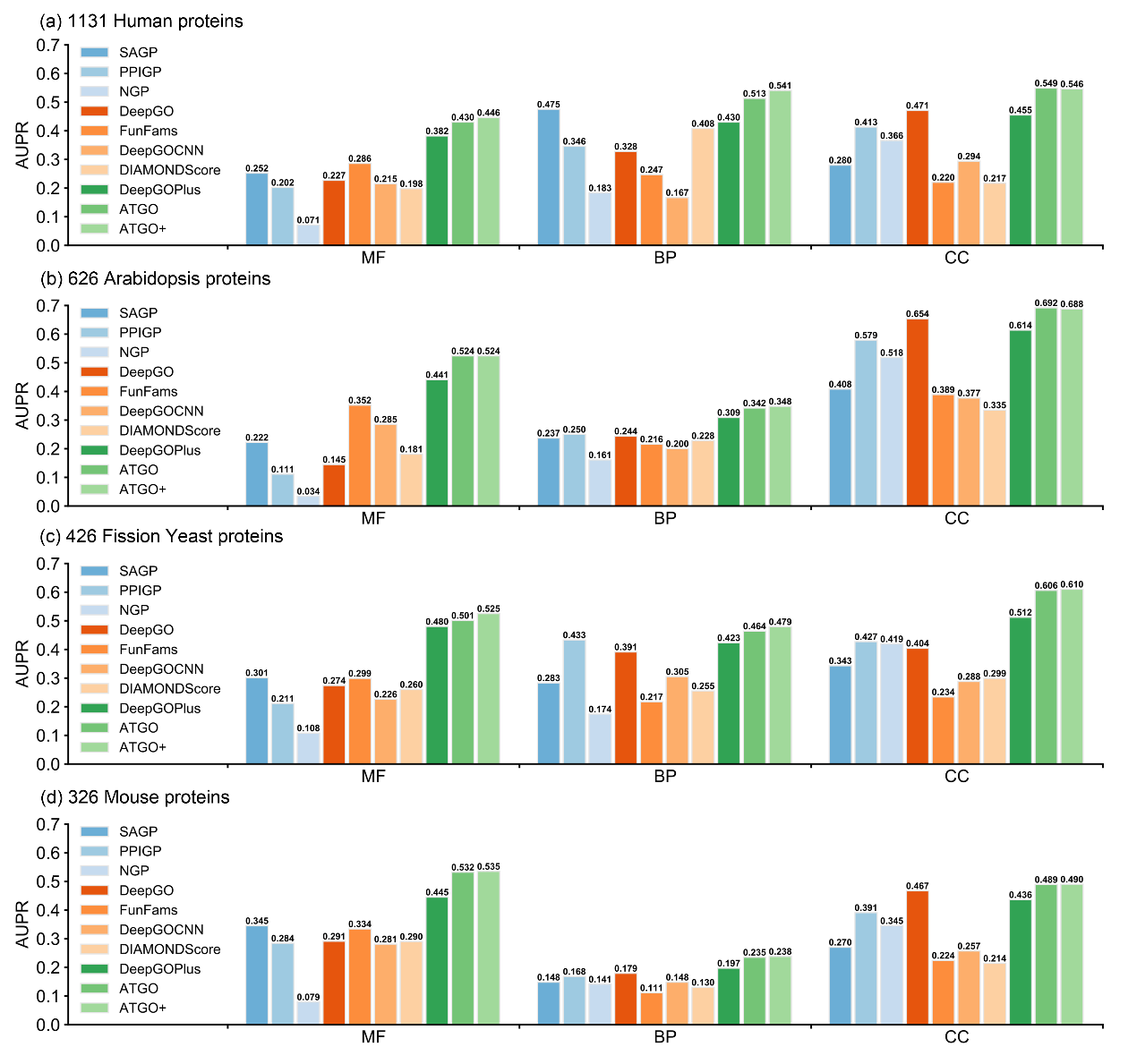

**Fig S4.** The AUPR values of ten GO prediction methods under the sequence identity cut-off $t_{1}=30\%$ for three GO aspects on four individual species in CAFA3 test dataset. **(a)** Human **(b)** Arabidopsis **(c)** Fission Yeast **(d)** Mouse.

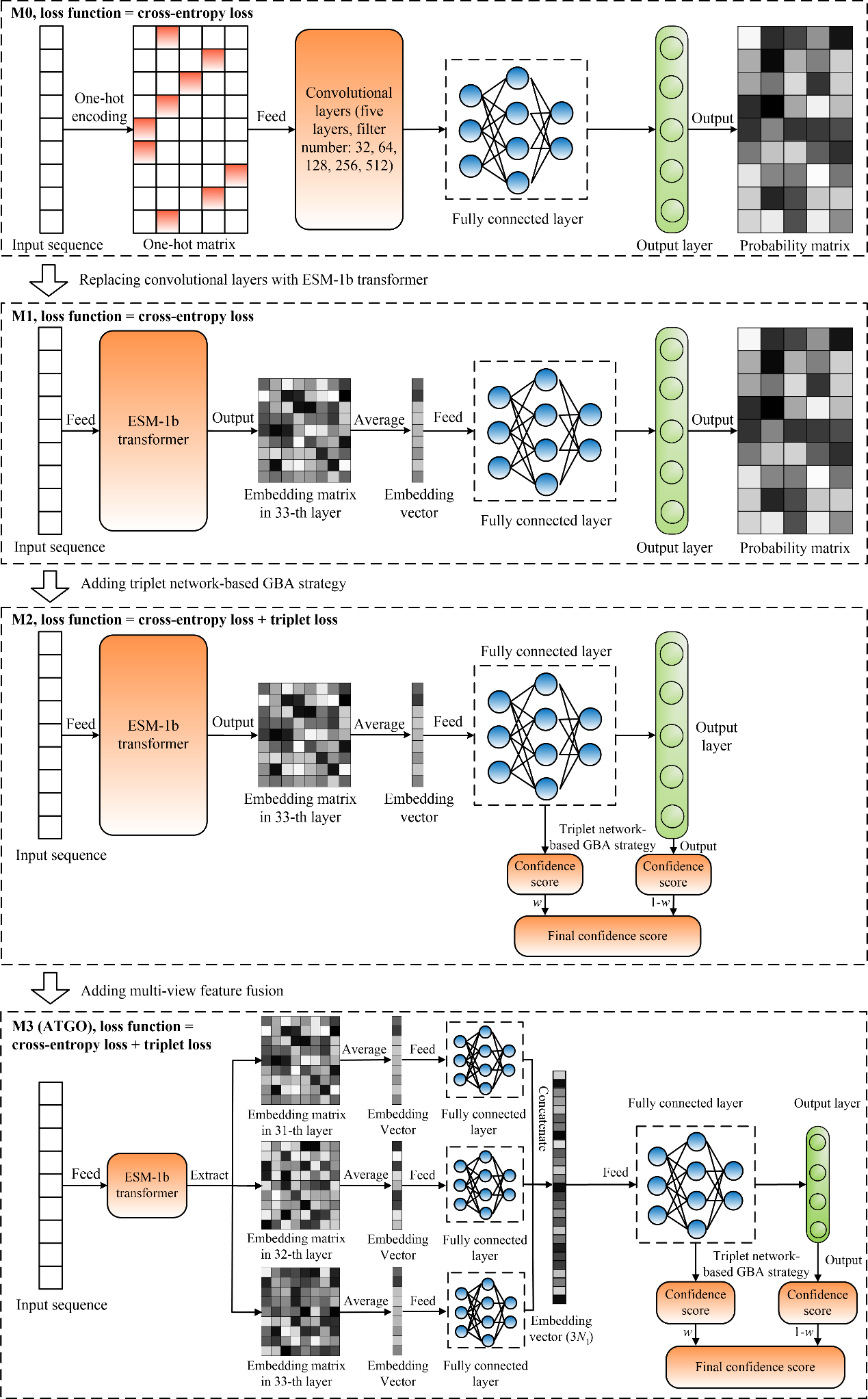

**Fig S5.** The architectures of different models in ablation study.

**Supporting Tables**

**Table S1**. The prediction performance with including root GO terms for ATGO and TALE on all 1068 test proteins. *p*-values in parenthesis are calculated between ATGO and TALE by one-tailed Student’s t-test. Bold fonts highlight the best performer in each category.

| **Method** | **Fmax** | | | **AUPR** | | |
| --- | --- | --- | --- | --- | --- | --- |
|  | **MF** | **BP** | **CC** | **MF** | **BP** | **CC** |
| TALE | 0.549  (7.3e-16) | 0.361  (2.4e-16) | 0.600  (1.1e-16) | 0.383  (2.6e-18) | 0.254  (1.4e-18) | 0.438  (4.7e-17) |
| ATGO | **0.688** | **0.465** | **0.695** | **0.689** | **0.413** | **0.654** |

**Table S2.** The summary of the proposed ATGO/ATGO+ and other ten competing GO prediction methods on a subset of 562 test proteins which have available templates or interaction partners in all of SAGP, PPIGP, FunFams, and DIAMONDScore. *p*-values in parenthesis are calculated between ATGO and other single-based methods and between ATGO+ and other composite methods by one-tailed Student’s t-test. Bold fonts highlight the best performer in each category.

| **Methods** | | **Fmax** | | | **AUPR** | | |
| --- | --- | --- | --- | --- | --- | --- | --- |
|  |  | **MF** | **BP** | **CC** | **MF** | **BP** | **CC** |
| Single algorithms | SAGP | 0.637 | 0.418 | 0.598 | 0.412 | 0.274 | 0.422 |
|  |  | (1.2e-06) | (4.6e-08) | (1.9e-10) | (5.2e-17) | (2.0e-15) | (8.5e-19) |
|  | PPIGP | 0.332 | 0.387 | 0.590 | 0.209 | 0.294 | 0.537 |
|  |  | (2.9e-17) | (1.2e-12) | (4.1e-11) | (1.5e-19) | (1.9e-14) | (1.4e-15) |
|  | NGP | 0.246 | 0.272 | 0.525 | 0.121 | 0.174 | 0.404 |
|  |  | (3.5e-18) | (1.2e-17) | (2.6e-14) | (2.7e-20) | (2.3e-18) | (4.0e-19) |
|  | DeepGO | 0.383 | 0.347 | 0.547 | 0.318 | 0.250 | 0.495 |
|  |  | (1.3e-16) | (3.9e-15) | (1.8e-13) | (2.1e-18) | (2.3e-16) | (4.5e-17) |
|  | FunFams | 0.512 | 0.343 | 0.482 | 0.325 | 0.176 | 0.293 |
|  |  | (3.9e-14) | (2.4e-15) | (1.5e-15) | (2.6e-18) | (2.5e-18) | (1.0e-20) |
|  | DeepGOCNN | 0.352 | 0.309 | 0.494 | 0.282 | 0.212 | 0.366 |
|  |  | (5.2e-17) | (1.4e-16) | (3.1e-15) | (8.1e-19) | (1.7e-17) | (9.6e-20) |
|  | DIAMONDScore | 0.629 | 0.405 | 0.580 | 0.322 | 0.232 | 0.316 |
|  |  | (7.3e-08) | (1.1e-10) | (8.6e-12) | (2.4e-18) | (6.1e-17) | (2.0e-20) |
|  | TALE | 0.397 | 0.318 | 0.527 | 0.338 | 0.243 | 0.499 |
|  |  | (2.2e-16) | (2.7e-16) | (3.0e-14) | (3.9e-18) | (1.3e-16) | (6.0e-17) |
|  | ATGO | 0.662 | 0.439 | 0.645 | 0.647 | 0.373 | 0.633 |
| Composite algorithms | DeepGOPlus | 0.641 | 0.412 | 0.580 | 0.581 | 0.335 | 0.543 |
|  |  | (9.8e-09) | (2.0e-10) | (6.9e-14) | (2.4e-13) | (4.5e-16) | (1.3e-16) |
|  | TALE+ | 0.640 | 0.423 | 0.611 | 0.588 | 0.346 | 0.621 |
|  |  | (7.1e-09) | (1.2e-08) | (1.7e-11) | (6.5e-13) | (6.4e-15) | (5.1e-10) |
|  | ATGO+ | **0.666** | **0.445** | **0.648** | **0.651** | **0.383** | **0.643** |

**Table S3.** The performance of ten GO prediction methods under the cut-off $t_{1}=30\%$ on 1177 no-knowledge (NK) and 2151 limited-knowledge (LK) CAFA3 proteins. Bold fonts highlight the best performer in each category.

| **Dataset** | **Method** | **Fmax** | | | **AUPR** | | | **Coverage** | | |
| --- | --- | --- | --- | --- | --- | --- | --- | --- | --- | --- |
|  |  | **MF** | **BP** | **CC** | **MF** | **BP** | **CC** | **MF** | **BP** | **CC** |
| NK proteins | SAGP | 0.467 | 0.351 | 0.472 | 0.248 | 0.189 | 0.280 | 0.84 | 0.85 | 0.83 |
|  | PPIGP | 0.286 | 0.316 | 0.476 | 0.176 | 0.212 | 0.440 | 0.85 | 0.83 | 0.84 |
|  | NGP | 0.184 | 0.260 | 0.467 | 0.083 | 0.159 | 0.380 | 1.00 | 1.00 | 1.00 |
|  | DeepGO | 0.302 | 0.332 | 0.502 | 0.230 | 0.233 | 0.501 | 1.00 | 1.00 | 1.00 |
|  | FunFams | 0.461 | 0.356 | 0.430 | 0.282 | 0.181 | 0.250 | 0.63 | 0.63 | 0.61 |
|  | DeepGOCNN | 0.267 | 0.304 | 0.428 | 0.203 | 0.193 | 0.297 | 1.00 | 1.00 | 1.00 |
|  | DIAMONDScore | 0.463 | 0.350 | 0.462 | 0.196 | 0.171 | 0.225 | 0.78 | 0.79 | 0.78 |
|  | ATGO | 0.513 | 0.393 | 0.557 | 0.472 | 0.314 | **0.559** | 1.00 | 1.00 | 1.00 |
|  | DeepGOPlus | 0.473 | 0.373 | 0.473 | 0.385 | 0.269 | 0.472 | 1.00 | 1.00 | 1.00 |
|  | ATGO+ | **0.523** | **0.396** | **0.557** | **0.482** | **0.316** | 0.555 | 1.00 | 1.00 | 1.00 |
| LK proteins | SAGP | 0.461 | 0.548 | 0.479 | 0.241 | 0.392 | 0.322 | 0.81 | 0.93 | 0.88 |
|  | PPIGP | 0.224 | 0.422 | 0.421 | 0.138 | 0.357 | 0.394 | 0.92 | 0.92 | 0.83 |
|  | NGP | 0.142 | 0.339 | 0.416 | 0.055 | 0.175 | 0.348 | 1.00 | 1.00 | 1.00 |
|  | DeepGO | 0.259 | 0.423 | 0.469 | 0.176 | 0.333 | 0.468 | 1.00 | 1.00 | 1.00 |
|  | FunFams | 0.481 | 0.472 | 0.508 | 0.320 | 0.243 | 0.332 | 0.67 | 0.76 | 0.75 |
|  | DeepGOCNN | 0.342 | 0.284 | 0.392 | 0.251 | 0.191 | 0.275 | 1.00 | 1.00 | 1.00 |
|  | DIAMONDScore | 0.452 | 0.518 | 0.469 | 0.200 | 0.344 | 0.256 | 0.74 | 0.89 | 0.84 |
|  | ATGO | 0.498 | 0.564 | 0.523 | 0.468 | 0.465 | 0.528 | 1.00 | 1.00 | 1.00 |
|  | DeepGOPlus | 0.449 | 0.524 | 0.478 | 0.394 | 0.401 | 0.470 | 1.00 | 1.00 | 1.00 |
|  | ATGO+ | **0.511** | **0.574** | **0.525** | **0.473** | **0.488** | **0.534** | 1.00 | 1.00 | 1.00 |

**Table S4.** The numbers of proteins for 20 species in CAFA3 test dataset.

| **Species name** | **Taxonomy ID** | **Sample number** |
| --- | --- | --- |
| Human | 9606 | 1131 |
| Arabidopsis | 3702 | 626 |
| Fission Yeast | 284812 | 426 |
| Mouse | 10090 | 326 |
| Escherichia Coli | 83333 | 224 |
| Fly | 7227 | 209 |
| Rat | 10116 | 97 |
| Bacillus Subtilis | 224308 | 76 |
| Dictyostelium Discoideum | 44689 | 49 |
| Zebrafish | 7955 | 46 |
| Budding Yeast | 559292 | 32 |
| Candida Albicans | 237561 | 27 |
| Salmonella Enterica | 99287 | 16 |
| Xenopus Laevis | 8355 | 14 |
| Methanocaldococcus Jannaschii | 243232 | 7 |
| Pseudomonas Putida | 160488 | 7 |
| Helicobacter Pylori | 85962 | 5 |
| Saccharolobus Solfataricus P2 | 273057 | 4 |
| Mycoplasma Genitalium | 243273 | 3 |
| Pseudomonas Aeruginosa | 208963 | 3 |

**Table S5.** The prediction performance of five GO prediction methods under the cut-off $t_{1}=100\%$ on CAFA3 test proteins. Bold fonts highlight the best performer in each category.

| **Dataset** | **Method** | **Fmax** | | | **AUPR** | | |
| --- | --- | --- | --- | --- | --- | --- | --- |
|  |  | **MF** | **BP** | **CC** | **MF** | **BP** | **CC** |
| All 3328 proteins | SAGP | 0.520 | 0.515 | 0.504 | 0.328 | 0.366 | 0.350 |
|  | PPIGP | 0.253 | 0.390 | 0.473 | 0.160 | 0.312 | 0.461 |
|  | NGP | 0.166 | 0.302 | 0.445 | 0.065 | 0.170 | 0.366 |
|  | ATGO | 0.548 | 0.520 | 0.555 | 0.504 | 0.445 | 0.551 |
|  | ATGO+ | **0.551** | **0.540** | **0.559** | **0.514** | **0.470** | **0.546** |
| 1177 no-knowledge proteins | SAGP | 0.494 | 0.387 | 0.509 | 0.297 | 0.230 | 0.337 |
|  | PPIGP | 0.299 | 0.335 | 0.491 | 0.189 | 0.236 | 0.471 |
|  | NGP | 0.192 | 0.260 | 0.467 | 0.082 | 0.160 | 0.380 |
|  | ATGO | **0.533** | 0.400 | 0.569 | 0.476 | 0.338 | 0.562 |
|  | ATGO+ | 0.532 | **0.419** | **0.570** | **0.490** | **0.346** | **0.553** |
| 2151 limited-knowledge proteins | SAGP | 0.537 | 0.602 | 0.498 | 0.347 | 0.473 | 0.366 |
|  | PPIGP | 0.221 | 0.435 | 0.456 | 0.140 | 0.366 | 0.447 |
|  | NGP | 0.147 | 0.339 | 0.416 | 0.055 | 0.175 | 0.348 |
|  | ATGO | 0.562 | 0.602 | 0.539 | 0.526 | 0.529 | 0.535 |
|  | ATGO+ | **0.566** | **0.622** | **0.542** | **0.532** | **0.564** | **0.537** |

**Table S6.** The incorrectly predicted GO terms for twelve methods on three proteins in BP aspect.

| **Method** | **A6XMY0** | **E7CIP7** | **F4I082** |
| --- | --- | --- | --- |
| SAGP |  | GO:0044419 GO:0009607 GO:0009605 GO:0043207 GO:0050896 GO:0009617 GO:0006952 GO:0006950 GO:0051707 | GO:0032502 GO:0042335 GO:0006869 GO:0006810 GO:0071702 GO:0051234 GO:0051179 GO:0048856 |
| PPIGP |  |  | GO:0032502 GO:0009628 GO:0044238 GO:0009987 GO:0044237 GO:0071704 GO:0009058 GO:0006807 GO:0065007 GO:0016043 GO:0008152 GO:1901576 GO:0042221 GO:0044249 GO:0050789 GO:0071840 |
| NGP | GO:0032502 GO:0044238 GO:0019222 GO:0060255 GO:0048856 GO:0044237 GO:0050794 GO:0071704 GO:0006807 GO:0065007 GO:0016043 GO:0008152 GO:0048518 GO:0043170 GO:0050789 GO:0071840 | GO:0032502 GO:0048856 GO:0019222 GO:0060255 GO:0050896 GO:0009987 GO:0044237 GO:0050794 GO:0006807 GO:0065007 GO:0016043 GO:0048518 GO:0050789 GO:0071840 | GO:0032502 GO:0044238 GO:0048856 GO:0019222 GO:0060255 GO:0009987 GO:0044237 GO:0050794 GO:0071704 GO:0006807 GO:0065007 GO:0016043 GO:0008152 GO:0048518 GO:0043170 GO:0071840 GO:0050789 |
| DeepGO | GO:0080090 GO:0019222 GO:0031326 GO:0031323 GO:0050789 GO:0071704 GO:2000112 GO:0060255 GO:0065007 GO:0048518 GO:0048519 GO:0065008 GO:0010468 GO:0019219 GO:0048583 GO:0009889 GO:1903506 GO:0050794 GO:0051171 GO:0008152 GO:2001141 GO:0051704 GO:0044238 GO:0051252 GO:0044237 GO:0051239 GO:0010556 GO:0048523 GO:0048522 | GO:2001141 GO:0009987 GO:0080090 GO:0019222 GO:0019219 GO:0009889 GO:0050896 GO:0051252 GO:0031323 GO:1903506 GO:0050794 GO:0006355 GO:0010556 GO:0065007 GO:0051171 GO:0031326 GO:0060255 GO:0010468 GO:2000112 GO:0050789 | GO:0048856 GO:0009892 GO:0080090 GO:0019222 GO:0009891 GO:0031327 GO:0031326 GO:0031325 GO:2000241 GO:0031323 GO:0048580 GO:0010629 GO:0032501 GO:0007165 GO:0050789 GO:0009893 GO:0009755 GO:0010605 GO:0009890 GO:0033993 GO:0031328 GO:2000112 GO:2000113 GO:0009737 GO:0060255 GO:0065007 GO:0048518 GO:0048519 GO:0010468 GO:0032502 GO:0009719 GO:0031324 GO:0048608 GO:0050793 GO:0009987 GO:0009889 GO:1903506 GO:0050794 GO:0003006 GO:0001101 GO:0051239 GO:2001141 GO:0042221 GO:0010033 GO:1901700 GO:0022414 GO:0009725 GO:0051252 GO:0019219 GO:0006355 GO:0010556 GO:2000026 GO:0048522 GO:0097305 GO:0010558 GO:0048523 GO:0051171 |
| FunFams |  | GO:0002376 GO:0009607 GO:0009605 GO:0043207 GO:0050896 GO:0009617 GO:0006952 GO:0006950 GO:0098542 GO:0042742 GO:0044419 GO:0050829 GO:0051707 GO:0006955 |  |
| DeepGOCNN | GO:0048583 GO:0023052 GO:0007165 GO:0023051 GO:0050829 GO:0010646 GO:0050789 GO:0051716 GO:0009966 GO:0051179 GO:0065007 GO:0065009 GO:0003008 GO:0007186 GO:0032502 GO:0032501 GO:0006810 GO:0050794 GO:0050830 GO:0007267 GO:0007154 GO:0051234 GO:0055085 GO:0051704 GO:0010469 GO:0048856 | GO:0032501 GO:0009605 GO:0050896 GO:0009987 GO:0071554 GO:0006950 GO:0065007 GO:0051179 GO:0051704 | GO:0044238 GO:0009987 GO:0044237 GO:0050794 GO:0071704 GO:0065007 GO:0016043 GO:0008152 GO:0051179 GO:0050789 GO:0071840 |
| DIAMONDScore |  | GO:0044419 GO:0009607 GO:0009605 GO:0043207 GO:0050896 GO:0009617 GO:0006952 GO:0006950 GO:0051707 |  |
| TALE | GO:0050829 GO:0032501 GO:0065007 GO:0050789 GO:0050794 | GO:0071554 GO:0009987 | GO:0044238 GO:0009987 GO:0044237 GO:0050794 GO:0071704 GO:0006807 GO:0065007 GO:0008152 GO:0050789 |
| ATGO |  | GO:0044419 GO:0009987 | GO:0032502 GO:0009628 GO:0003006 GO:0009987 GO:0022414 GO:0065007 GO:0050789 |
| DeepGOPlus | GO:0032501 GO:0065007 | GO:0044419 GO:0009617 GO:0009607 GO:0009605 GO:0043207 GO:0050896 GO:0009987 GO:0006952 GO:0006950 GO:0051707 | GO:0009987 |
| TALE+ |  | GO:0009617 GO:0009607 GO:0009605 GO:0043207 GO:0050896 GO:0009987 GO:0044419 GO:0006950 GO:0051707 | GO:0009987 |
| ATGO+ |  | GO:0044419 GO:0050896 | GO:0032502 GO:0042335 GO:0048856 |

**Table S7.** The numbers of proteins and GO terms in benchmark dataset.

| **Benchmark dataset** | $\boldsymbol{N}_{\boldsymbol{MF}}^{\boldsymbol{P}}$ | $\boldsymbol{N}_{\boldsymbol{BP}}^{\boldsymbol{P}}$ | $\boldsymbol{N}_{\boldsymbol{CC}}^{\boldsymbol{P}}$ | $\boldsymbol{N}_{\boldsymbol{ALL}}^{\boldsymbol{P}}$ | $\boldsymbol{N}_{\boldsymbol{MF}}^{\boldsymbol{T}}$ | $\boldsymbol{N}_{\boldsymbol{BP}}^{\boldsymbol{T}}$ | $\boldsymbol{N}_{\boldsymbol{CC}}^{\boldsymbol{T}}$ | $\boldsymbol{N}_{\boldsymbol{ALL}}^{\boldsymbol{T}}$ |
| --- | --- | --- | --- | --- | --- | --- | --- | --- |
| Training dataset | 49135 | 79491 | 71982 | 109132 | 6581 | 20882 | 2782 | 30245 |
| Validation dataset | 515 | 860 | 664 | 1089 | 818 | 3894 | 417 | 5129 |
| Test dataset | 577 | 839 | 586 | 1068 | 876 | 3469 | 382 | 4727 |

$N_{MF}^{P}$/$N_{BP}^{P}$/$N_{CC}^{P}$/$N_{ALL}^{P}$: The number of proteins for MF/BP/CC/all three aspects.

$N_{MF}^{T}$/$N_{BP}^{T}$/$N_{CC}^{T}$/$N_{ALL}^{T}$: The number of GO terms for MF/BP/CC/all three aspects.

**Table S8.** The values of $margin$, $c_{f}$, $\alpha$, and $K$ for three GO aspects

| **GO aspect** | $\boldsymbol{margin}$ | $\boldsymbol{c}_{\boldsymbol{f}}$ | $\boldsymbol{\alpha}$ | $\boldsymbol{K}$ |
| --- | --- | --- | --- | --- |
| MF | 0.1 | 0.8 | 5 | 30 |
| BP | 0.1 | 0.8 | 5 | 100 |
| CC | 0.1 | 0.8 | 5 | 100 |

**Supporting Texts**

**Text S1. Sequence alignment-based GO prediction (SAGP)**

In SAGP, we select the function templates, which share high sequence similarity with the query, to annotate its function. Specifically, for a query sequence, Blast software (1) is used to scan the corresponding templates with an e-value cutoff of 0.1. The confidence score of the GO term $q$ by SAGP is calculated by

${S\left( q \right)}_{SAGP}=\frac{\sum_{k=1}^{n} b_{k}\cdot I_{k}(q)}{\sum_{k=1}^{n} b_{k}}$	(S1)

where $n$ is the number of templates identified, $b_{k}$ is the bit-score of $k$-th template by Blast; $I_{k}\left( q \right)=1$, if the $k$-th template is associated with $q$ in the experimental function annotation; otherwise, $I_{k}\left( q \right)=0$.

**Text S2. Protein-protein interaction-based GO prediction (PPIGP)**

For a query, we search its interaction partners from the STRING database (2) for functional annotation. Then, we remove the interaction partners which are not found in the training dataset. Finally, the remaining partners are used to annotate the query. The confidence score is calculated using the same scoring function as in SAGP (i.e., Eq. S1), where $b_{k}$ is the score assigned by STRING as confidence of interaction between the query and the $k$-th partner.

**Text S3. Naïve-based GO prediction (NGP)**

In NGP, the confidence score that a query is associated with GO term $q$ is calculated by the frequency of $q$ in the training dataset:

${S\left( q \right)}_{NGP}=N(q)/N_{GO}$	(S2)

where $N(q)$ is the number of proteins associated with $q$, and $N_{GO}$ is the number of proteins with at least one annotation for the same GO aspect as $q$. This predictor can be thought of as a prior arising from the overall abundance of a particular annotation in the training dataset.

**Text S4. The mathematics formulas for ESM-1b transformer**

The formulas for scale dot-product attention are described as follows

$A^{ij}=softmax(M_{Q}^{ij}{M_{K}^{ij}}^{T}/\sqrt{d_{ij}}) M_{V}^{ij}$	(S3)

$M_{Q}^{ij}=H^{i}W_{Q}^{ij}, M_{K}^{ij}=H^{i}W_{K}^{ij}, M_{V}^{ij}=H^{i}W_{V}^{ij}$	(S4)

where $A^{ij}$ is the attention matrix in the $i$-th layer and $j$-th head; $M_{Q}^{ij}$, $M_{K}^{ij}$, and $M_{V}^{ij}$ are Query, Key, and Value matrices in the $i$-th layer and $j$-th head, and $W_{Q}^{ij}$, $W_{K}^{ij}$, and $W_{V}^{ij}$ are the corresponding weight matrices, respectively; $H^{i}$ is the input matrix in the $i$-th layer (e.g., $H^{1}$ is the inputted one-hot coding matrix); and $d_{ij}$ is the scale parameter.

The outputs of all attention heads in $i$-th layer are concatenated as a new matrix, which is further fed to multilayer perceptron (MLP):

$A^{i}=A^{i1}A^{i2}\ldots A^{im}$	(S5)

$E^{i}=max\left( A^{i}W_{1}^{i}+b_{1}^{i}, 0 \right)W_{2}^{i}+b_{2}^{i}$	(S6)

where $A^{i}$ is the concatenated attention matrix in the $i$-th layer, $E^{i}$ is the feature embedding in the $i$-th layer, $W_{1}^{i}$ and $W_{2}^{i}$ are weight matrices, $b_{1}^{i}$ and $b_{2}^{i}$ are bias matrices.

The output of the last attention layer is fed to a fully connected layer with SoftMax function to generate a $L\times28$ probability matrix:

$P=SoftMax\left( E^{n}W^{n}+b^{n} \right)$	(S7)

where the ($l$-th, $c$-th) value in $P$ indicates the probability that the $l$-th token in the masked sequence is predicted as the $c$-th type of amino acid.

The loss function is designed as:

${Loss}_{esm}=E_{x\sim X}\sum_{l\in x(M)} \left( -\frac{logP_{l, c\left( l \right)}}{\left| x\left( M \right) \right|} \right)$	(S8)

where $x$ is a sequence in training protein set $X$, $x(M)$ is a set of masking position in $x$, $\left| x(M) \right|$ is the number of elements in $x(M)$, $c(l)$ is the type index of amino acid for the $l$-th token in $x$ before masking, and -$logP_{l, c\left( l \right)}$ is negative log likelihood of the true amino acid $x_{l}$ under condition of masking.

**Text S5. The functional similarity between two proteins**

The functional similarity of two proteins is measured by the F1-score between their GO terms:

$F1-score=2(pre\times rec)/(pre+rec)$, $pre=ns/n_{a}$, $rec=ns/n_{b}$	(S9)

where $ns$ is the number of same GO terms between two proteins, $n_{a}$ and $n_{b}$ are the numbers of GO terms for proteins $a$ and $b$, respectively.
